# Towards comprehensive and quantitative proteomics for diagnosis and therapy of human disease

**DOI:** 10.1101/072215

**Authors:** Paolo Cifani, Alex Kentsis

## Abstract

**Abbreviations:**

DDA: Data Dependent Acquisition
DIA: Data Independent Acquisition
PRM: Parallel Reaction Monitoring
PTM: post-translational modification
SAX: strong anion exchange (chromatography)
SCX: Strong cation exchange (chromatography)

**Abstract:** Despite superior analytical features, mass spectrometry proteomics remains seldom used for the basic investigation and clinical treatment of human disease. This need is particularly pressing for childhood diseases that can be rare in incidence and variable in presentation. Modern mass spectrometry enables detailed functional characterization of the pathogenic biochemical processes, as achieved by accurate and comprehensive quantification of proteins and their regulatory chemical modifications. Here, we describe how high-accuracy mass spectrometry in combination with high-resolution chromatographic separations can be leveraged to meet these analytical requirements in a mechanism-focused manner. We review the quantification methods capable of producing accurate measurements of protein abundance and post-translational modification stoichiometries. We then discuss how experimental design and chromatographic resolution can be leveraged to achieve comprehensive functional characterization of biochemical processes in complex biological proteomes. Finally, we describe current approaches for quantitative analysis of a common functional protein modification: reversible phosphorylation. In all, current instrumentation and methods of high-resolution chromatography and mass spectrometry proteomics are poised for immediate translation into improved diagnostic and therapeutic strategies for pediatric and adult diseases.

## Introduction

Ever since the first discovery of specific proteins associated with human disease [1], the field of protein chemistry and later proteomics sought to identify new and improved markers of disease and targets of therapies. While the instrumentation for analytical chemistry and mass spectrometry has steadily improved, incorporation of this approach into preclinical investigation and clinical care has lagged [2]. With notable exceptions, such as mass spectrometry-based detection of bacterial pathogens [3], and drug and metabolites [4,5], recent advances in mass spectrometry remain largely confined to analytical chemistry laboratories [6]. Recently, we and others have sought to apply high-accuracy mass spectrometry [7] approaches for the discovery of improved diagnostic markers and therapeutic targets [8-16]. As a result of these and other studies, several methodological requirements for translational and clinical proteomics have emerged, including the need to balance analytical sensitivity and accuracy with the breadth of analyte detection, as driven by sample throughput. Here, we review the recently developed mass spectrometric methods in their current ability to enable comprehensive and quantitative proteomics, as they relate to the translational and clinical applications.

## Biological Mass Spectrometry Proteomics

Protein activities in cells are controlled by multiple factors, including but not limited to protein synthesis and degradation [17], alternative splicing [18], post-translational chemical modification [19], intra-cellular localization [20], and interaction with co-factors and regulators [21]. Understanding differential regulation of all these mechanisms requires accurate quantification of proteins and their proteo- and chemoforms, which is increasingly being achieved by combining mass spectrometry-based proteomics with biochemical techniques and computational analyses [22-25]. These approaches generate data of increasing breadth and depth, as evidenced by the recently established workflows for mass spectrometric detection of post-translationally modified peptides [26, 27]. The general analytical requirement to obtain such biologically meaningful data is the need to accurately and sensitively measure the abundance of all relevant protein chemoforms in a sample. Here, we will focus on bottom-up proteomics approaches, which analyze peptides generated by enzymatic or chemical proteolysis instead of the corresponding intact proteins, as this approach remains the most prevalent today [7, 28], though recent improvements in intact protein analysis should lend themselves to large-scale intact proteomics [29].

## Quantitative Proteomics

High-throughput quantification of proteins and peptides historically relied on dye fluorescence intensity of gel resolved proteins, i.e., DIGE [30], or on correlative measures such as for example the number of fragmentation spectra recorded for a given protein [31]. Nowadays, these methods are used less frequently, because improvements in chromatography, ionization, mass spectrometry instrumentation, and data analysis enable more accurate quantification by direct measure of currents generated by specific peptide ions. The signal produced depends not only on the specific analyte concentration, but also on the efficiency of formation of the relative ions (ionization and fragmentation properties, as applicable). As a result, ion current-based quantification is always a relative and sample-specific measure.

With the exception of methods dependent on reporter ions, discussed later, quantification of peptides by mass spectrometry requires multiple measurements of the current generated by specific ions. These measurements are integrated in the time domain of the corresponding chromatographic peak to calculate the area under the curve (AUC), which is the complete quantitation metric [32, 33]. This method is more robust than instantaneous ion current measurements, reducing the variability produced by differential chromatographic properties of peptides and variable ionization efficiencies.

Using modern software, specific ion currents can be extracted from any series of mass spectra. For example, signal intensity of un-fragmented peptide ions can be retrieved from full-range high-resolution data-dependent precursor scans [32, 33], a strategy that in principle enables proteome-wide quantification. However, far higher sensitivity, precision, and linear dynamic range are achieved by targeted quantification, which consists of detecting a ions within defined *m*/*z* windows selected by mass filters of increasing resolving power. The most widespread implementation, still considered the gold-standard for peptide quantitation, is selected reaction monitoring (SRM, also referred to as MRM for multiple reaction monitoring), which uses triple-stage quadrupole instruments to first filter specific *m*/*z* range for fragmentation and subsequently filter specific fragment ions produced by collision-induced dissociation before dynode detection [34, 35]. This method benefits from the high sensitivity of dynode detectors, and the robustness conferred by the uninterrupted ion beam, but is limited by the relatively low resolution of current mass filters that hinders the specificity of the assays, which thus require careful validation [36, 37].

Parallel reaction monitoring (PRM) is conceptually similar to SRM in the use of mass filtering of narrow precursor isolation windows, but uses high-resolution mass analyzers, such as the Orbitrap, to enable acquisition of complete high-resolution fragment ion spectra [38, 39]. While comparable in sensitivity to SRM, PRM enables potentially complete sequencing of the target peptide, with the consequent improvements in specificity and accuracy of quantitation. However, its higher duty cycle may reduce assay multiplexing, a drawback recently alleviated by the introduction of the isotope-triggered PRM approaches [40]. Both methods enable absolute sensitivity in the attomolar range, and up to five order of magnitude of linear dynamic range, which is still less than the biologic concentration range of proteins in human tissues [41, 42].

On the other hand, data independent acquisition (DIA) in principle can overcome the limited throughput of targeted methods by iteratively selecting portions of the *m*/*z* range for fragmentation, prior to high-resolution detection of fragments from all the filtered precursor ions. Subsequent deconvolution of these fragmentation spectra permits peptide identification and extraction of chromatographic elution peaks for quantification [43-47]. While recent improvements in the resolution of time-of-flight spectrometers, such as the parallel accumulation-serial fragmentation (PASEF) method [48], promise to increase the instrumental duty cycle to permit data independent analysis of increasing sensitivity and accuracy, recent benchmarking of DIA using existing instruments demonstrated lower accuracy as compared to PRM and SRM [49].

An alternative strategy for peptide quantitation leverages the detection of reporter ions generated by the fragmentation of chemically reactive isobaric tags, such as for example iTRAQ and TMT [50, 51]. Both reagents consist of an isotopically encoded reporter ion, an amine reactive *N*-hydroxysuccinimidyl moiety, and a normalizing group to ensure that precursors labeled with different isotopologues remain isobaric are thus co-selected for fragmentation. These reagents are particularly useful in clinical applications as they enable isotopic labeling of samples derived from human tissues, but require controls for variable labeling efficiency and limited dynamic range [52].

## Towards Comprehensive Quantification

While current approaches for quantitative mass spectrometry are sufficiently accurate to permit robust peptide quantification, they have yet to be applied for comprehensive analyses. For example, a typical SRM assay with chromatographic scheduling can monitor on the order of 100 peptides. Conversely, DDA experiments, implementing either precursor ion current or reporter ion quantification, permit measuring the abundance of several thousand peptides across multiple samples, although with reduced precision, reproducibility and sensitivity. These observations provided the rationale to consider targeted approaches as a mere validation method for comprehensive DDA surveys. However, it is important to note that the complexity of mammalian tryptic proteomes far exceeds the sequencing duty cycle of current instruments [53], and that DDA is biased towards abundant and readily ionizable peptides that often do not include analytes of interest [54]. As a consequence, these approaches may not be suitable for the analysis of relevant molecular markers.

However, for many human diseases, including childhood diseases, comprehensive proteomic profiling may not be necessary, as relevant molecular markers have been identified using hypothesis-based or other high-throughput approaches such as genomics. For example, numerous childhood and adult cancers exhibit oncogenic activation of kinase signaling [55, 56], and chromatin and gene expression regulatory pathways [57, 58]. Thus, measurements of biologically or pathologically meaningful analytes may not require ‘whole-proteome’ approaches, and instead may rely on quantification of marker panels defined to probe specific pathways, as for example the PI3K-mTOR/MAPK signaling cascade [59] or the DNA damage response network [60]. This can also involve knowledge-based “sentinel” proteins [61], or other markers of pathway activity, such as those generated by reduced representation approaches [62]. Collections of SRM assays for this purpose have already begun development for cancer and infectious diseases [63-66].

The major determinant of throughput for both analytes and specimens is the duty cycle of targeted mass spectrometric detection in relation to the time scale of analytical chromatographic separation. One obvious solution for this problem involves enhancing chromatographic resolution prior to MS analysis to obtain adequate separation over extended chromatographic gradients [67]. This rationale was indeed successfully applied to increase the number of targeted mass spectrometry assays scheduled in a single experiment [68]. Improved chromatographic resolution can also be achieved by multi-dimensional and orthogonal separation techniques [69, 70], which also provide a means to improve mass spectral sampling, and detection and quantification of low abundance ions, thereby increasing the exposure of specific proteome subsets such as post-translationally modified peptides [7, 71-73]. However, most offline sample fractionation workflows are potentially hindered by sample losses that limit their overall robustness and reproducibility [74]. Online chromatographic fractionation has been successfully applied to DDA experiments, demonstrating high efficiency and sensitivity due to automation and reduced sample requirements [75-78]. In unpublished results from our laboratory, we observed that automated online fractionation using multi-dimensional chromatography efficiently and reproducibly separated peptides from low-abundance transcription factors from other abundant isobaric ions co-eluting in final chromatographic dimension coupled to nanoelectrospray ionization. This enabled accurate quantification by targeted precursor and fragment ion detection of analytes that were otherwise not detected at all using conventional offline multi-dimensional or online single dimensional chromatographic separations.

Due to the variability of peptide ionization and fragmentation, all quantitative methods based on ion current extraction are inherently relative in nature [32, 33, 79]. Extracted ion chromatograms can be matched to compare the signal produced by the same peptide in different experiments. Such label-free methods have been used for comprehensive analysis of phosphorylation stoichiometry in model cell systems [32], [33, 80]. This strategy was also used in translational and preclinical studies to identify human disease biomarkers [12, 14, 16]. However, far more accurate measurements can be achieved using synthetic external reference peptides by comparing the signals produced by isotopologue peptides undergoing simultaneous chromatographic separation and ionization, thus minimizing technical variability and noise. Such approaches require isotopically encoded reference peptides for all the targeted analytes. Metabolic labeling of cell lines or primary cells *in vitro* has been used to generate reference standards for relative quantification of tumor samples [81-83]. However, it is still unclear whether such standards sufficiently capture the complexity of biologically variable analytes, such as specific post-translational modifications. Moreover, differential protein turn-over rates may lead to uneven proteome labeling [17]. Tissue samples can also be directly labeled using isotopically encoded chemical reagents including cysteine reactive moieties [84], ^18^O water [85], iTRAQ and TMT reagents [50, 51] as well as other amine reactive groups producing dimethyl [86, 87] or nicotinic acid derivative [88, 89] adducts. While permitting universal labeling for quantitative mass spectrometry, such approaches require controls for variable or non-specific labeling. Alternatively, quantitation can be achieved using isotopologue synthetic peptides, as they can be introduced at known concentrations directly, thus enabling absolute quantification [35, 90].

## Towards Comprehensive Functional Proteomics

Along with protein abundance, measured by quantification of the corresponding peptides, post-translational protein modifications are biologically important regulatory mechanisms that currently can be analyzed best using quantitative mass spectrometry [91]. In particular, the well-established regulatory functions of protein kinase signaling led to the refinement of methods for enrichment and analysis of phosphorylated peptides. Mass spectrometry is particularly well suited for characterization of protein chemoforms, as specific chemical modifications produce specific diagnostic alterations of peptide molecular mass. However, the sub-stoichiometric nature of protein phosphorylation and the relatively low abundance of many kinases and kinase substrates pose serious challenges for robust measurements of site occupancies and stoichiometries. Instrumental advances that enable robust phospho-proteomics include the development of specific affinity chromatography reagents and chromatographic strategies for the enrichment of phosphorylated peptides [71-72, 92-96]. Such approaches, for example, have recently been used to measure biological kinetic processes [97], and have been successfully coupled to targeted detection for enhanced sensitivity [98].

Enrichment of phosphorylated peptides is most commonly achieved using offline separations, that despite efforts towards miniaturization and automation [62, 99], are still prone to variable adsorptive losses that can potentially confound quantification measurements. To overcome this limitation, online chromatographic enrichment of phosphorylated peptides has been developed [100, 101]. Importantly, the detection of phosphorylated peptides does not appear to be significantly affected by their intrinsic chromatographic and ionization properties [28], suggesting that improved exposure afforded by online multi-dimensional chromatography might enable robust and sensitive quantitative analysis. Consistent with this notion, enhanced detection of phosphorylated peptides was observed using online fractionation by combining alkaline reverse phase and strong-anion exchange chromatography [76, 77]. Importantly, these automated multi-dimensional chromatographic methods might improve the detection and quantitation of other chemically modified, e.g., acetylated, methylated etc, peptides without the need for dedicated affinity enrichment procedures, thus providing a generalized method for quantitative functional proteomics [71].

## Future Directions

There is a clear and unmet need for improved strategies for diagnosis, prognostication, and treatment of human. Current and emerging methods for high-resolution chromatography and mass spectrometry now enable routine accurate and sensitive quantitation of many biologically and pathologically relevant biomarkers. In particular, modern mass spectrometry satisfies the analytical requirements for comprehensive functional proteomics. Targeted bottom-up proteomics enable accurate quantification over a wide range of analyte concentrations present in clinical tissue specimens. In addition to data independent approaches, recent advances in mechanism-based analysis of specific cellular processes may permit clinically relevant quantification of biologically or pathologically functional proteome subsets. Specifically, this is empowered by robust and reproducible sample processing and fractionation, which is now achievable using automated online multidimensional chromatography systems. This should enable not only precision functional proteomics by improving targeted detection of chemically modified peptides and proteins, but also provide specific mechanistic information into biological and disease processes themselves.

## Acknowledgements

We thank John Philip for comments on the manuscript. This work was supported by the American-Italian Cancer Foundation (P.C.), NIH R21 CA188881, P30 CA008748, Alex’s Lemonade Stand Foundation, Gabrielle’s Angel Foundation, and the Damon Runyon-Richard Lumsden Foundation Clinical Investigator Program (A.K.).

